# Oxygen-induced stress reveals context-specific gene regulatory effects in human brain organoids

**DOI:** 10.1101/2024.09.03.611030

**Authors:** Benjamin D. Umans, Yoav Gilad

## Abstract

The interaction between genetic variants and environmental stressors is key to understanding the mechanisms underlying neurological diseases. In this study, we used human brain organoids to explore how varying oxygen levels expose context-dependent gene regulatory effects. By subjecting a genetically diverse panel of 21 brain organoids to hypoxic and hyperoxic conditions, we identified thousands of gene regulatory changes that are undetectable under baseline conditions, with 1,745 trait-associated genes showing regulatory effects only in response to oxygen stress. To capture more nuanced transcriptional patterns, we employed topic modeling, which revealed context-specific gene regulation linked to dynamic cellular processes and environmental responses, offering a deeper understanding of how gene regulation is modulated in the brain. These findings underscore the importance of genotype-environment interactions in genetic studies of neurological disorders and provide new insights into the hidden regulatory mechanisms influenced by environmental factors in the brain.

## Introduction

Understanding how gene regulatory variants function across different cellular and environmental contexts is essential for interpreting genetic associations with disease. Gene-by-environment (GxE) interactions occur when genetic variants influence how individuals respond to specific environmental exposures, leading to inter-individual differences in phenotypes, including variability in disease susceptibility. This concept is particularly important in complex diseases, where individuals with different genetic backgrounds may exhibit varying risk profiles for conditions such as neuropsychiatric disorders [1,2]. For instance, environmental factors such as stress [3–5], oxygen deprivation [6,7], or infection [8–13] can trigger disease-relevant gene regulatory effects that remain hidden in static, steady-state conditions.

Gene regulatory catalogs like GTEx (Genotype-Tissue Expression project [14]) provide valuable insights into how genetic variants affect gene expression across various tissues in steady-state conditions. However, the majority of disease-associated loci remain unexplained, likely due to their regulatory effects being specific to certain cell types or environmental contexts that have not been fully explored [15,16]. This gap is particularly pronounced in the brain, where the complex interplay between different cell types and environmental stressors can contribute to the onset and progression of neurological and psychiatric diseases [17–22].

Brain cells, especially neurons, are highly sensitive to environmental perturbations like hypoxia (oxygen deprivation), given the brain’s high metabolic demand and susceptibility to oxidative damage [23]. Hypoxia is a well-known neurological risk factor throughout life, arising from conditions such as sleep apnea, high altitude, respiratory infections, and premature birth [24–28]. Hypoxic exposure has profound effects on cognitive function, white matter integrity, and increase the risk for neurodegeneration and psychiatric disorders [29–47]. Despite the importance of oxygen homeostasis and hypoxia to brain function, we lack comprehensive insight into how different brain cell types respond to these environmental stressors at the gene regulatory level, which limits our ability to interpret genetic associations with neurological traits [47–50].

In this study, we used human brain organoids to investigate the transcriptional responses of diverse brain cell types to oxygen perturbation across 21 individuals. By applying single-cell RNA sequencing, we captured gene expression data both under baseline conditions and after exposure to varying oxygen levels. Through whole-genome sequencing of each donor, we identified genetic contributions to these responses, revealing dynamic gene regulatory effects with significant relevance to neurological and psychiatric disease susceptibility. This approach goes beyond characterizing gene regulatory variation in static, post-mortem tissue and opens new avenues for studying GxE interactions in a controlled, *in vitro* setting.

## Results

We differentiated brain organoids from the iPSCs of 21 unrelated Yoruba individuals from Ibadan, Nigeria [51] (see Methods). We performed oxygen manipulation experiments in two batches of 7-16 individuals, with two individuals replicated across batches to allow us to control and account for batch effects (**Figure 1a**). Specifically, following seven weeks of growth at atmospheric oxygen levels (21% O_2_), organoids were adapted to 10% O_2_ to mimic the physiologic environment experienced by brain cells *in vivo*. After one week of culture at 10% O_2_, organoids were either maintained at physiologic oxygen (baseline/normoxia), transferred to low oxygen (1% O_2_; hypoxia), or transferred to high oxygen (21% O_2_; hyperoxia) for 24 hours. Following the treatments, we dissociated organoids in the presence of transcriptional inhibitors and multiplexed equal proportions of each sample in preparation for single-cell RNA-sequencing, targeting 3,000 cells per individual and oxygen condition, and a depth of 20,000 reads per cell. After demultiplexing and quality control, we retained data from 170,841 cells (normoxia: 52,671, hypoxia: 57,788, hyperoxia: 60,382; median 5,666 UMI counts per cell).

**Figure 1.**
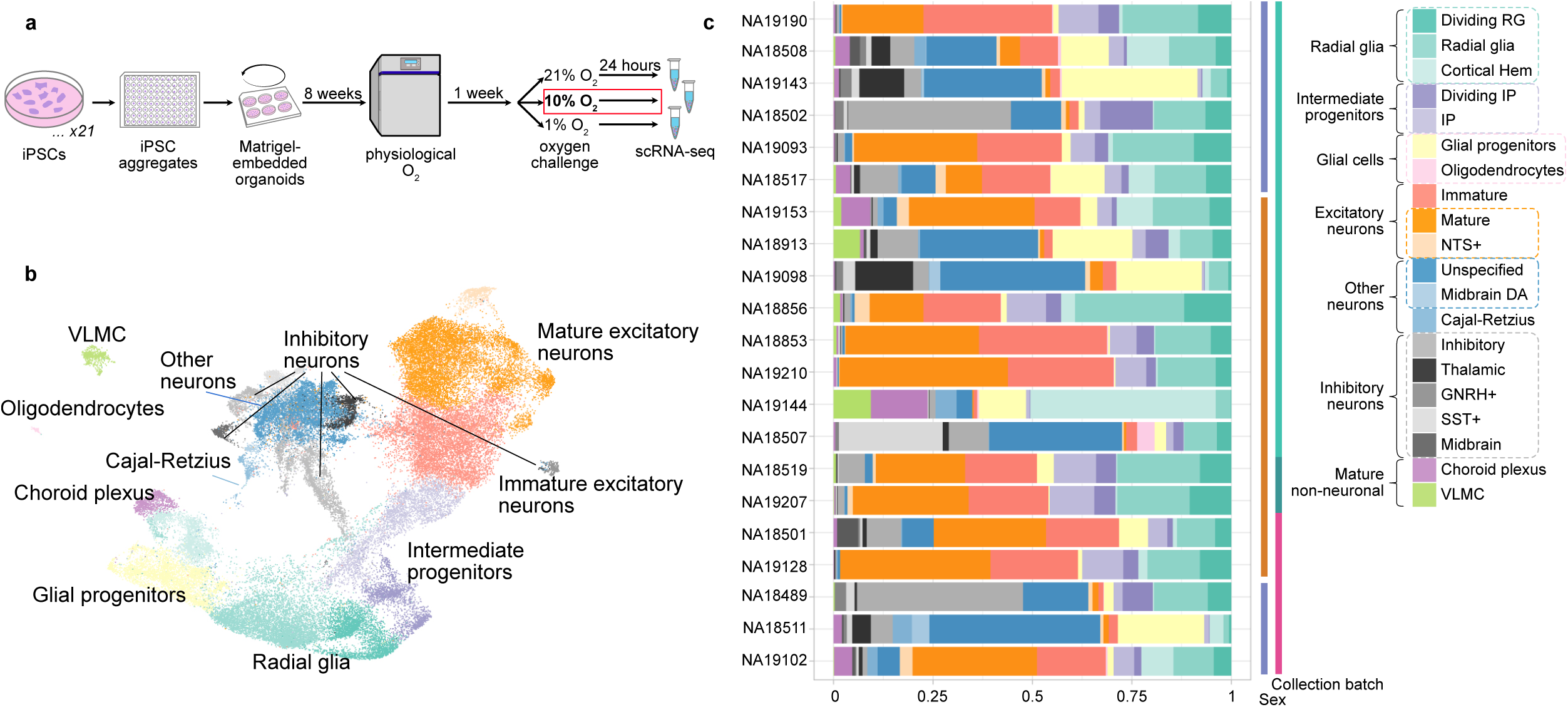
A panel of brain organoids yields diverse cell types across individuals. **(a)** Workflow of data collection. Brain organoids were first differentiated (see Methods) and adapted to physiological oxygen before 24-hour exposure to high, low, or control oxygen levels. Data shown in this figure are taken from the 10% “normoxia” condition. **(b)** UMAP representation of organoid single-cell transcriptomes highlighting principal cell types obtained across all individuals in our iPSC panel in the normoxia condition. **(c)** Proportion of cells from each parental cell line by annotation, with colors corresponding to the UMAP shown in (b).

### Brain organoids comprise diverse cortical and non-cortical cell types

We first sought to characterize the cell type composition of brain organoids maintained at baseline oxygen levels. We annotated cell clusters using fetal brain reference data and known marker genes [52,53], finding a variety of cortical cell types, including radial glial progenitors, intermediate progenitors, excitatory neurons, and inhibitory neurons (**Figure 1b, S1**, **Methods**). We also identified a substantial cluster of neurons with non-cortical identities, including thalamic and midbrain inhibitory neurons and GNRH^+^ cells (**Figure 1b, S1,** Methods). Out of 20 high-confidence cell types, 10 are present in over half of the individuals, with a median of 11 cell types detected per individual (**Figure 1c**).

We used *propeller* [54] to assess differences in cell type proportions across treatment conditions, and found that organoid composition was largely unaltered by oxygen manipulation (**Figure S2a, Table S1**). Moreover, the cell type composition of an additional set of control organoids which we maintained at atmospheric oxygen levels for the duration of the experiment did not differ substantially from what we observed in the treatment conditions (**Figure S1b**). We observed differences in cell type composition across individuals, with the rarer cell types (such as midbrain dopaminergic-like cells, mature oligodendrocytes, vascular leptomeningeal cells, and Cajal-Retzius cells) present in only a minority of samples (**Figure 1, S1e**). However, across treatments, the cell type composition of organoids from the same individuals generally remained similar (with the exception of NA19144; **Figure S2a**), suggesting that the treatments did not have a marked effect on cellular composition.

Since the treatments did not seem to result in noticeable differences in cell composition, we focused again on the differences in cell composition across individuals. We found no effect of sex or iPSC passage number on cell type proportions, but we observed that certain rare cell types – including oligodendrocytes, inhibitory neuron subtypes (midbrain, thalamic, and SST^+^), and midbrain dopaminergic-like cells – differed in proportion across individuals from different collection batches (**Table S1**). To assess the extent of confounding by batch, we merged biologically similar clusters to generate a set of 10 coarse annotations representing the principal cell types in our data (**Figure S1d, Methods**). When we examined cell type proportions among coarsely-defined clusters, the batch effect disappeared, suggesting that principal cortical cell types were largely stable across experiments. Still, we performed all subsequent analyses using both annotations to assess potential bias caused by over clustering (**Methods**), and to ensure that the two approaches produce similar outcomes. In what follows, we report findings from the more interpretable fine-grained annotation set (results from the coarse clustering approach are provided in **Figure S1d, S1e, S2d-e, S4a, S4b, S5b-f**).

### Shared transcriptional response patterns reveal cell-type-specific vulnerabilities to hypoxia

To identify differentially expressed (DE) genes between baseline and either hypoxic or hyperoxic conditions, we applied a linear mixed model to pseudobulk expression data for each cell type and treatment condition[55]. We identified a total of 10,230 DE genes in response to hypoxia, ranging from 91 to 5,590 genes per cell type (FDR<0.05, **S Table S2**). Similarly, we identified 10,425 hyperoxia-responsive genes, ranging from 17 to 6,102 genes per cell type (FDR<0.05, **S Table S2**). In at least one (any) cell type, 2,703 hypoxia-responsive genes (and correspondingly, 2,855 hyperoxia-responsive genes) exhibited a greater than 1.5-fold change in expression compared to baseline. As expected, we detected far more DE genes in abundant cell types (**Figure S2c**), within which 76-92% of hypoxia-evoked transcriptional effects and 84-93% of hyperoxia-evoked transcriptional effects were modest (smaller than 1.5-fold).

We were intrigued by the large differences in the numbers of DE genes across cell types, an observation that cannot be fully explained by cell abundance (Figure S3). To explore this further, we analyzed the data using multivariate adaptive shrinkage (*mash*) [56], to account for incomplete power and assess the similarity of transcriptional responses across treatment conditions and cell types. By combining power across cell types, we were able to detect weak DE effects that emerge in multiple cell types and, importantly, accurately identify condition- and cell-type-specific DE genes.

As expected, we observed similar oxygen response patterns among related cell types (**Figure 2a**). Even using *mash*, we found that more than half of the oxygen-responsive genes we identified (68% of hypoxia-responsive genes and 63% of hyperoxia-responsive genes, FDR<0.05, effect size >1.5-fold) were DE in fewer than three cell types, consistent with the idea that oxygen homeostasis mechanisms are tuned to the needs of distinct brain cell types [57]. We also observed a tendency for DE genes with large effects to be more cell-type-specific than DE genes with smaller effects (**Figure S2b**), an observation that is counter-intuitive with respect to power considerations.

**Figure 2.**
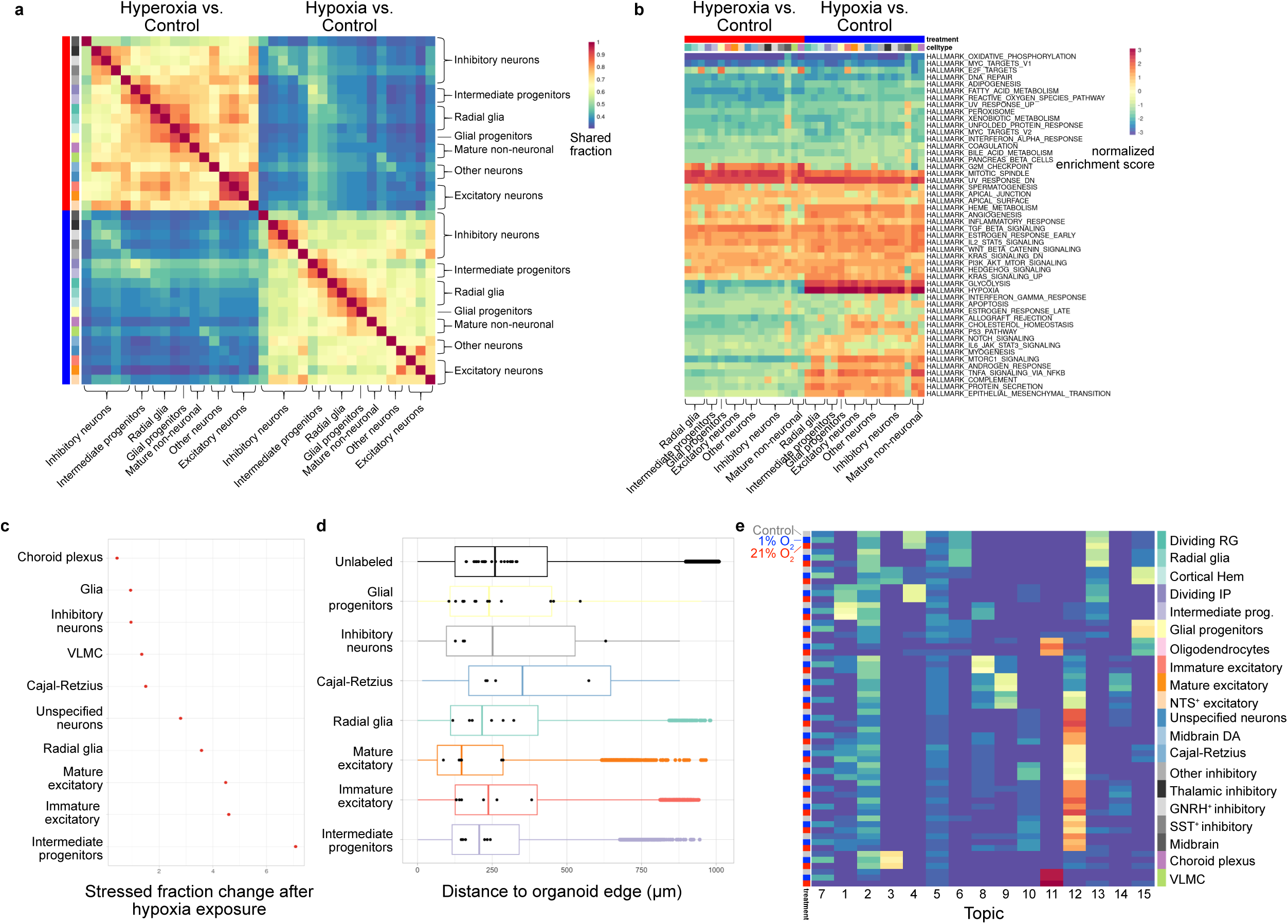
Transcriptional responses of brain organoids to oxygen changes. **(a)** Fraction of shared differential expression effects between cell types and conditions. Sharing was assessed from MASH posterior estimates of significance and effect sizes (see Methods and Figure S2). **(b)** Enrichment of gene annotations among differentially expressed genes across cell types and conditions. Annotations are taken from the MSigDB Hallmark gene sets, with enrichments calculated by *fgsea*. **(c)** Fractional change in hypoxia-stressed fraction of each cell type after hypoxia exposure (coarse cell classification). Cells were classified as hypoxia-stressed or hyperoxia-stressed by *Gruffi* using a gene set derived from treatment responses shared across all cell types. See also Figure S4. **(d)** Distribution of cell distances to organoid perimeters measured after immunofluorescent labeling. “Unlabeled” measurements are derived from DAPI-labeled nuclei with no immunofluorescent label. Black dots represent sample means. Note that “Dividing Cells” encompass both dividing radial glia and dividing intermediate progenitors. **(e)** Average topic loading for each cell type and treatment condition. Topic 7 tracks hypoxia exposure, while other topics reflect processes found in one or several cell types.

Most cell types exhibited a modest positive correlation between the transcriptional responses to hypoxia and hyperoxia (**Figure 2a, S2**), and many gene families related to general response to stress were enriched among DE genes in both treatment conditions. For example, all oxygen-treated cell types are enriched for DE genes with roles in cellular metabolism and inflammation, and most are enriched for DE genes involved in cell proliferation and apoptosis (**Figure 2b**). Reassuringly, hypoxia, specifically, also induced a well-recognized core regulatory program across all cell types, including the upregulation of hypoxia-inducible factor (HIF) target genes [58–60].

As our observations pointed to a general stress response following the treatments, we sought to characterize the proportion of single cells expressing regulatory signatures of stress in baseline and treatment conditions, so we can specifically focus on the response to the treatments. To parse cell type heterogeneity in the transcriptional response to oxygen perturbation, and to differentiate between stressed and unstressed cells, we identified gene sets that capture robust and general responses to each treatment condition (**Table S3, Methods**). We leveraged these gene sets in a granular filtering approach [61] to classify cells as either stressed or unstressed, and then repeated our differential expression analysis after censoring the stressed cells. We found that although stressed cells contribute disproportionately to the treatment expression response, they do not account for it entirely: in cell types with a higher proportion (12-36%) of stress-censored cells, the number of DE genes (FDR<0.05, fold-change>1.5) decreased by as much as 64% relative to randomly censored data (**Figure S4d**).

Cell types that include an increased fraction of stressed cells following exposure to hypoxia or hyperoxia may be especially sensitive to oxygen perturbation. We calculated the change in the proportion of stressed cells between baseline and treatment conditions to assess cell-type-specific sensitivity within brain organoids. We found that intermediate progenitors, immature neurons, and radial glia are particularly responsive to both hypoxia and hyperoxia (**Figure 2c, S4**).

The observation of large differences in cell-type sensitivity to changes in oxygen may be significant and help us better understand disease mechanisms. It is possible, however, that cell-type-specific sensitivity to the treatments could be explained by variation in organoid spatial structure. To examine this, we used antibody markers to map eight major cell types in cryosections obtained from the same organoids we used for sequencing (**Methods**). We then quantified the accessibility of each cell type to exogenous oxygen by measuring the distance from immunostained cells to the organoid periphery and compared the distance distribution across cell types. While different cell types were distributed at various depths within each organoid, variation between organoids was comparable to the differences observed between cell types (**Figure 2d**; ANOVA F-test of organoid-level medians, p=0.342). On the whole, the most oxygen-sensitive cell types were localized neither more superficially nor more deeply than other cell types. Thus, cell type-specific sensitivity to oxygen perturbation appears to be driven primarily by cellular identity, rather than cell position within the organoids.

### Context-specific responses to oxygen perturbation

We observed that oxygen perturbation induced widespread transcriptional effects, many of which were shared among subsets of cells within and between cell types. Perturbed cells continued to express cell-type-specific markers, simultaneously maintaining their respective identities while adopting a more general signature of oxidative stress (**Figure S4c**). This suggests that discrete cell types may fail to capture continuous contextual responses, including signatures shared by developmentally related or physically proximal cells. In an effort to capture the subtler transcriptional patterns in our data, we used topic modeling to decompose cellular transcriptomes into 15 groups (topics) that capture distinct sources of transcriptional variation (**Methods**).

Several topics largely recapitulate discrete cell type classifications, as expected, consistent with the notion that cell identity is the primary source of transcriptional heterogeneity in our data (**Figure 2e**). Other topics, such as topic 7, recapitulate properties that were already known to us, such as the collection of DE genes that we previously used to identify hypoxia-stressed cells (**Figure S3d**). We also identified topics that captured dynamic cellular processes and developmental states. Still, several topics revealed new patterns: topic 4 is shared by dividing radial glia and dividing intermediate progenitors and is distinct from separate topics that tightly correlate with each of these cell identities in isolation. Indeed, topic 4 is defined by elevated expression of genes involved in DNA replication and cell division, including *MKI67, TOP2A*, and centrosomal proteins (**Figure S3d**), capturing shared aspects of the dividing cell environment across different cell types. In turn, topic 3 shows modest loading in cortical hem progenitors and higher loading in choroid plexus cells, reflecting their shared developmental origins: signals from the cortical hem influence the differentiation and patterning of cells at the boundary of the cerebral cortex and hippocampus, including those forming the choroid plexus [62]. Altogether, topic modeling allowed us to recapture functional relationships and shared states that were concealed in our analysis of discrete, mutually-exclusive cell type clusters, providing us with an enriched view of brain organoid dynamics.

### Transcriptional responses to oxygen perturbation are genetically regulated

Having detected thousands of genes that are differentially expressed in response to oxygen perturbation, we sought to uncover potential genetic sources for inter-individual variation in treatment response. We aggregated single-cell gene expression data into pseudobulk groups defined by their unique combination of donor, cell type, and treatment condition. After excluding groups comprising fewer than 20 cells, we removed cell types that had fewer than seven individuals in each treatment condition (**Methods**). For each of the 14 remaining cell types, we separately mapped *cis* eQTLs under hypoxic, hyperoxic, and baseline conditions, including gene expression principal components as model covariates to account for the effects of sex, batch, and latent confounding factors (**Methods**). We expected most eQTLs to be shared across treatment conditions – indeed, our DE analysis indicates that most genes are robust to oxygen perturbation. We therefore used *mash* to weigh evidence for SNP-gene associations across all three treatment conditions, considering each cell type in turn. As we and others have found, this approach improves power to detect individually weak signals that emerge consistently in multiple experimental contexts [63–65].

Across 14 cell types, we tested a total of 9,478 genes and found 36,778 *cis* eQTLs in 8,320 genes, with a median of four eQTLs per eGene (local false sign rate < 0.05). Among these, we identified 14,358 standard eQTLs (in 5,952 eGenes), in which the eQTL effect size is of similar size and direction across all treatment conditions (we used a conservative 2.5-fold cutoff to define similar effect sizes across treatment conditions, as we consider shared effects to be the null; **Table S4**). We also identified 22,420 oxygen-response eQTLs in 7,338 genes, namely eQTLs that have significantly different effect sizes between treatment conditions. Note that the sum of eGenes associated with at least one standard eQTL and those associated with at least one oxygen-responsive eQTLs is larger than 8,320, because eGenes are often associated with more than a single eQTL across cell types, and often with both standard and eQTL oxygen-responsive eQTLs in different cell types Oxygen-responsive eQTLs included 3,687 loci associated with distinct effects in the hypoxia condition (in at least one cell type), 3,603 loci associated with distinct effects in hyperoxia, and 2,935 loci associated with a difference between the baseline normoxia and both treatments (**Figure 3a**). Of particular note, across cell types, 15,045 oxygen response eQTLs are not associated with a statistically significant eQTL in the baseline (normoxia) condition. The genetic effect of these loci on gene regulation in the different cell types can only be detected under the stress conditions imposed by the change in oxygen levels. Consistent with this, oxygen-responsive eQTLs – particularly those not found under normoxia – are associated with the expression of genes that are less likely to have eQTL effects in cerebral cortex tissues in GTEx (one-sided paired Wilcoxon test, P = 0.007; **Figure 3c, Figure S5a**).

**Figure 3.**
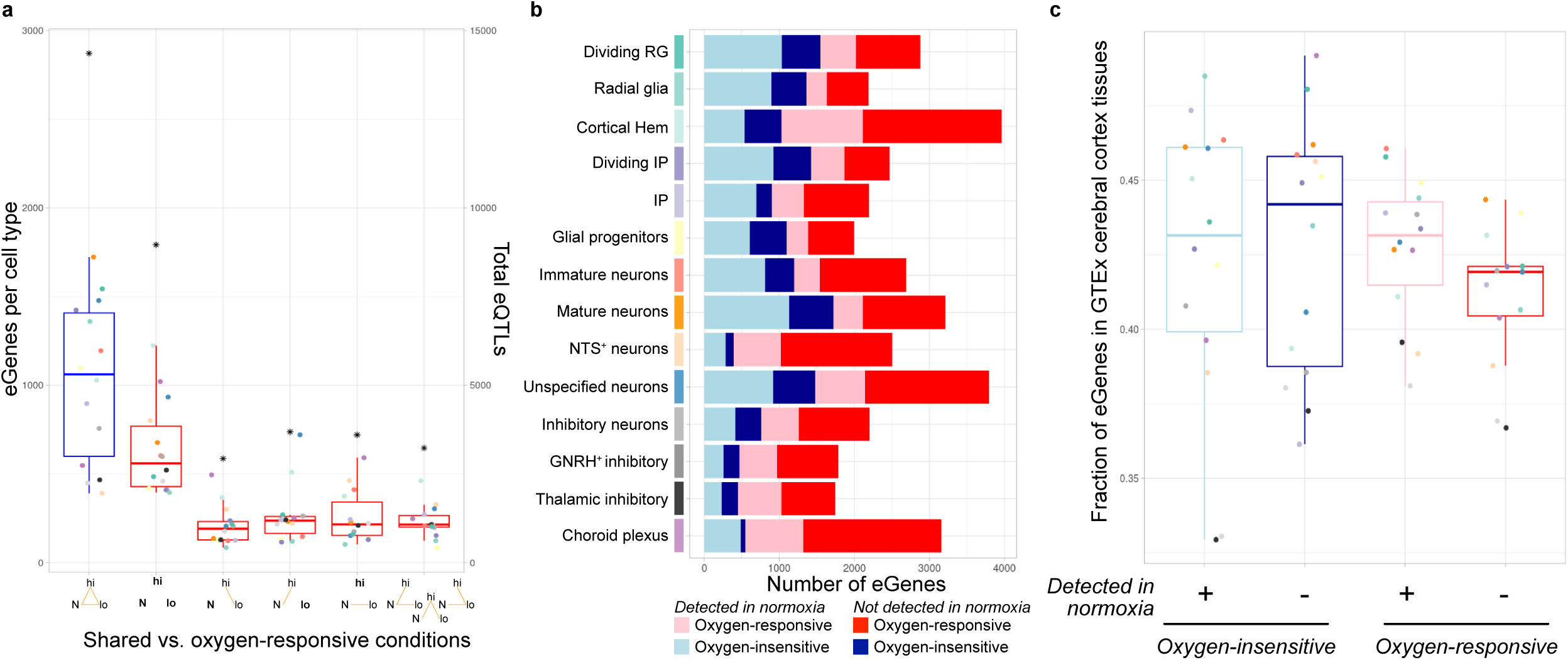
Discovery of treatment context-specific eQTL effects across cell types. **(a)** Number of eGenes identified in each cell type, classified as “standard” (blue) or oxygen-responsive (red) according to contexts in which effect sizes differ by less than 2.5-fold. For each category of oxygen-responsive eGenes, shared effects are indicated by orange lines and the condition with a uniquely different eQTL effect is bolded (N = normoxia, lo = hypoxia, hi = hyperoxia). Total eQTLs of each category are indicated by stars (right axis). **(b)** Number of eGenes identified in each cell type, classified by sharing across oxygen treatment conditions and detection in normoxia condition. Light and dark shades of red and blue categories, respectively, sum to the blue and red categories shown in (a). **(c)** Fraction of eGenes identified in this study that are classified as eGenes in GTEx cerebral cortex tissues (see Methods). Treatment context-specific eGenes that were not detected in control conditions were less likely to be found in GTEx (p=0.0067, one-sided paired Wilcoxon test of oxygen-insensitive category against oxygen-responsive/undetected in normoxia category).

### Context-dependent genetic regulation

In the analysis of gene expression levels, topic modeling allowed us to place cells along continuous axes of variation that are not predicated upon marker gene or reference annotations. We reasoned that topics could also be deployed to identify genetic regulatory effects that emerge in contexts defined more precisely than is possible using the discrete categories of cell type and treatment. To explore the effects of *cis* eQTLs in an expanded set of precisely-defined cellular contexts, we tested for interactions between eQTLs and topics. Rather than individually testing each eQTL-topic interaction, we used CellRegMap [66] to jointly test all linear combinations of topics, improving our power to detect a wide range of genotype-context interactions.

We identified 289 genes with a topic-interacting eQTL. To infer the relevant cellular context for each eQTL, we assessed the correlation between its estimated effect and the loading for each topic. When possible, we checked to ensure our interpretation was corroborated by results from our analysis of discrete cell types and treatment conditions. For example, topic 15 describes cortical hem and glial progenitor cells. Out of the top 12 eGenes associated with topic 15 (Pearson correlation > 0.6), 10 were identified as eGenes exclusively in cortical hem or glial progenitor cells using our pseudobulk approach. Also, among the top 12 eGenes is the cholesterol transporter *ABCA1* (r = 0.77), which showed modest eQTL effects in our pseudobulk analyses of cortical hem cells, glial progenitors, radial glia, and intermediate progenitors (**Figure 4a**). Of note, radial glia are the precursors to both glial progenitors and intermediate progenitors. Concordantly, our topic-based approach revealed a modest correlation between *ABCA1* and topic 6 (r=0.26), which is defined primarily by radial glia.

**Figure 4.**
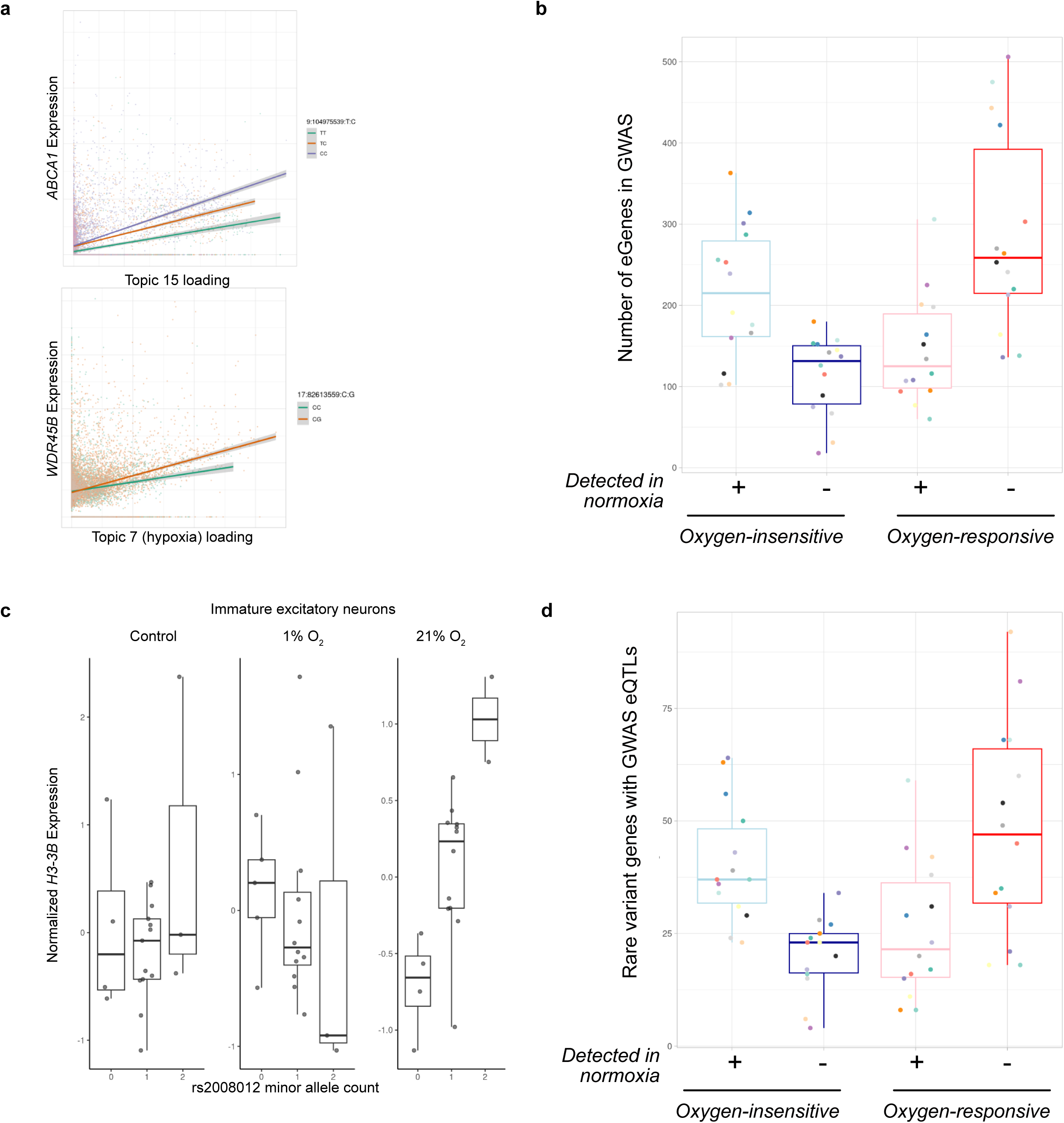
Organoid eQTLs can help interpret human disease genetics. **(a)** Example topic-interacting eQTLs. *ABCA1* expression is correlated with the inferred cell environment, as defined by linear combinations of topics, and this effect is largely driven by cortical hem and glial progenitor topic 15 in a genotype-dependent manner. The *WDR45B* eQTL effect is largely explained by hypoxia-associated topic 7. Each point corresponds to a single pseudocell used for CellRegMap topic interaction QTL mapping. **(b)** Number of eGenes in each cell type which are the nearest gene to a genome-wide significant GWAS finding among 402 brain-related traits. **(c)** Example of a cell type- and treatment-specific regulatory association matching a significant GWAS variant. **(d)** Number of eGenes in each cell type and discovery condition for which rare loss-of-function alleles have been associated with disease (see Methods).

Using the topic eQTL analysis we found 218 eGenes whose regulatory effects were significantly correlated with topic 7, which is associated with hypoxic stress (p<0.05, Bonferroni correction). Unexpectedly, 118 of these eGenes did not have hypoxia-specific eQTLs in our standard cell-type specific analysis, including *WDR45B* and *CD44*, which were more strongly correlated with topic 7 than any other topic. This is an example of the additional insight we can gain by using topics instead of discrete cell identities. *WDR45B* is a member of the WIPI protein family of autophagosomal proteins, and *WDR45B* mutations have been linked to numerous severe neurodevelopmental disorders [67]. *CD44* is a known regulatory target of HIF1 and interacts with HIF2 to modify local tissue responses to hypoxia [68,69]. While we identified multiple eQTLs for *WDR45B* using the standard analysis, they were either oxygen-insensitive or responsive specifically to *hyper*oxia. We also identified eQTLs for *CD44* that were oxygen-responsive, but not to hypoxia specifically. Taken together, these results highlight the utility of decomposing complex cellular states into constituent programs for the analysis of gene-by-environment interactions.

### Context-dependent eQTLs help to interpret the effects of disease-associated loci

The use of brain organoids allowed us to identify context-specific gene regulatory effects that are underrepresented in standard eQTL studies of post-mortem tissues (**Figure 3c, Figure S5a**). We reasoned that oxygen-responsive eQTLs could help to explain uncharacterized genetic associations with disease – particularly if the eQTL effects were undetectable at baseline. To explore this possibility, we examined the overlap of eGenes with GWAS loci assembled from 402 brain-relevant traits in each cell type (**Methods**). Across cell types, eGenes with response eQTLs that were latent at baseline (median 258.5 per cell type, 1,745 total) included a comparable number of disease-associated genes as standard eGenes detected at baseline (median 215 per cell type, 1,411 total; two-sided paired Wilcoxon test p=0.194; **Figure 4b**). Focusing on the 4,713 novel eGenes that were not represented in GTEx cerebral cortex tissues, we found an average of 158 disease-associated eGenes per cell type (total 1,014) to overlap oxygen-response eQTLs that were latent at baseline. Finally, of the 218 eGenes that interact with hypoxia (i.e., topic 7), 55 correspond to a GWAS gene, including 31 genes that are not eGenes in GTEx cortical tissue. Thus, mapping eQTLs in oxygen-treated brain organoids allowed us to uncover novel, disease-relevant effects that could not be detected in primary cortical tissues.

Next, we asked whether context-specific eQTLs could revise previous interpretation of disease-associated SNPs. Focusing on oxygen-responsive lead eQTL SNPs that were associated with brain traits in GWAS, we identified 146 associations (corresponding to 76 genes) in which eQTL mapping implicated a different target gene than the original GWAS report or a simple nearest-gene heuristic. For example, we identified a hyperoxia-specific association between rs2008012 and the expression of *H3F3B* in immature excitatory neurons and an oxygen-insensitive effect in dividing intermediate progenitors (**Figure 4c, Figure S5g**). rs2008012 is associated with variation in uncinate fasciculus white matter, which connects the limbic system to the brain’s frontal lobes [70]. This SNP has been described as an eQTL for ten different genes, principally in blood, with brain data pointing to effects on expression of *TRIM47, TRIM65, WBP2,* and *ACOX1* [71–73]. While both coding and regulatory mutations in *H3F3B* are known to cause severe neurodevelopmental phenotypes, our data suggest that subtle, context-specific regulation of *H3F3B* expression during development may also contribute to microstructural brain features.

Most genes have been associated with at least one regulatory eQTL. However, protein-coding mutations are relatively rare. We reasoned that genes harboring rare deleterious coding mutations might also be regulated by common variants – albeit with subtler phenotypic effects. To determine whether context-specific eQTLs can be used to connect common GWAS variants with rare disease-causing mutations, we assembled results from five large exome studies of neurological or psychiatric traits, finding 1,672 genes with rare variants that are associated with at least one disease or developmental condition (**Methods**). Out of these 1,672 genes, 905 are eGenes in at least one cell type or condition in our data, and 349 have a significant GWAS association (**Figure 4d**). We identified 37 cases (corresponding to 22 genes) in which the lead SNP was significantly associated with a brain-related GWAS trait (**Table S5**). For example, damaging missense mutations in *ATP2A2* confer risk for bipolar disorder (OR 10.4) [74]. In subtypes of radial glia and mature neurons, we identified a novel eQTL for *ATP2A2* (<u>rs4766428</u>) that is strongly associated with cognitive ability, risk of schizophrenia, and risk of anorexia nervosa [75–78]. Similarly, a radial glial eQTL (<u>rs9611486</u>) for *EP300*—rare variants of which have been associated with autism, developmental delay, and Rubinstein-Taybi syndrome 2—is a risk SNP for anxiety symptoms [79,80]. Importantly, nearly half of the associations found in this comparison were not detected under baseline oxygen conditions, highlighting the importance of examining diverse cell types and perturbed states in order to identify trait-relevant regulatory effects.

## Discussion

In this study, we measured cell-type-specific responses to oxygen stress in a genetically diverse panel of brain organoids. Oxygen perturbation induced a robust transcriptional response across all assayed cell types, which included a common oxygen stress response signature as well as cell-type-specific changes. Leveraging the genetic and cellular heterogeneity present in our organoid panel, we identified thousands of dynamic, oxygen-responsive eQTLs, many of which have effects that are undetectable at baseline and therefore are absent in data collected from post-mortem cortical tissues. Moreover, the use of topic modeling allowed us to identify genetic effects that transcend categorical notions of cellular identity to regulate cell division, differentiation, and other continuous processes. By collecting functional data from biological contexts that are difficult to access *in vivo*, we were able to characterize the putative regulatory role of hundreds of GWAS loci, many of which had never been associated with an eQTL. Our results show that repurposing organoids for gene-environment interaction studies is a valuable and increasingly tractable approach that complements population genomic studies of primary tissues, and may be particularly valuable for studies of neurological and psychiatric diseases.

Our organoid differentiation approach included a small molecule cocktail designed to promote dorsal telencephalic patterning, which we selected to mitigate the between-organoid variability previously observed in un-patterned cortical organoids [81]. We nonetheless found substantial differences among organoids derived from different donor cell lines, with some cell types present in only a minority of samples. As expected, this cell type heterogeneity resulted in incomplete power to map eQTLs across cell types, reflecting the trade-off between cell type resolution and abundance inherent to single-cell data analysis. At the same time, this heterogeneity allowed us to measure gene regulation in biological contexts that have never before been examined at the population level.

Though all cell types in our study responded to hypoxia, intermediate progenitor cells were among the most sensitive to hypoxic challenge. Interestingly, immature neurons showed increased expression of the intermediate progenitor cell marker *TBR2*/*EOMES* following 24-hour hypoxia exposure, possibly indicating a rapid transition from intermediate progenitor to neuronal identity. Paşca et al. demonstrated that a 48-hour hypoxic challenge led to an unfolded protein response (UPR)-dependent depletion of TBR2^+^ intermediate progenitors, which underwent an apparently precocious developmental transition to CTIP2^+^ neurons [82]. Although we observed minimal induction of transcriptional markers of UPR and found only modest changes in the overall abundance of intermediate progenitor cells, our results are consistent with these findings, and suggest that the effects of hypoxia on intermediate progenitor cell development are visible even after a shorter exposure period. It is also possible that a sustained 48-hour shift from 21% to <1% oxygen induces a stronger effect with greater dependence on the UPR than our paradigm, which includes a period of adaptation to physiologic oxygen and a shorter hypoxic treatment. In either case, our data suggest that intermediate progenitor cells are especially labile in the face of fluctuating environmental oxygen, and may play an outsized role in linking transient oxygen stress to brain phenotypes [48,82].

We characterized gene regulatory variation in the context of normoxia, hypoxia, and hyperoxia. Though acute hypoxia-induced brain injury is relatively rare, transient fluctuations in brain oxygen availability are common, suggesting that the gene regulatory effects in this study may be pervasive in the population. For example, sleep apnea, in which repeated episodes of oxygen desaturation and restoration conspire to produce persistent oxidative stress, is estimated to affect nearly a billion people worldwide [83]. Recent studies using *in vivo* oxygen biosensors have also observed regions of local tissue hypoxia in rodent brains, both at rest and during demanding tasks [84,85]. While the significance of these minute-to-minute environmental fluctuations are not yet clear, their existence raises the possibility that hypoxia-evoked transcriptional changes may be ongoing features of the brain under normal conditions. Our results imply that these conditions elicit a host of gene expression changes with varying consequences across the population, and that those differences may in turn affect complex brain-related traits.

Our study focused on just three experimental conditions. By extending the topic modeling framework to a wider range of treatments, future studies could determine how many of the dynamic eQTLs we discovered can also be identified in the presence of other *in vitro* stressors. Recent technical advances, including supplementing organoids with non-neuronal cell types [86] and *in vivo* implantation [87–89], promise to dramatically expand the scope of disease-relevant interactions that can be captured in brain organoids and further extend the approach employed here. We expect that future studies of regulatory variation in these contexts will help prioritize targets for *in vivo* experimental manipulation.

## Methods

### Stem cell culture and organoid formation

We generated brain organoids using 21 iPSC lines (12 male, 9 female) that belong to an extensively characterized panel of iPSCs derived from Yoruba individuals from Ibadan, Nigeria (YRI) [51]. Stem cells were maintained on Matrigel-coated plates and fed with StemFlex media (Gibco) supplemented with penicillin and streptomycin. Cells were passaged at least twice before organoid formation using 0.5 mM EDTA in PBS and seeded on new plates in the presence of CEPT [90].

Organoids were formed using a protocol modified from published methods [81,91–93]. Cells were dissociated using 0.5 mM EDTA in PBS and passed through a 40 µm filter, then aggregated by centrifugation in 96-well ultralow attachment round-bottom plates (Nunclon), with 10,000 cells per well in 100 µL of StemFlex medium with penicillin/streptomycin, 5 µM XAV, and CEPT. After 16-hour overnight incubation, medium was replaced with E6 medium supplemented with 100 nM LDN193189 (Cayman), 5 µM XAV939 (Cayman), 10 µM SB431542 (Cayman), 1X MEM non-essential amino acids (Gibco), and penicillin/streptomycin. Cell aggregates were fed with this medium every other day for seven days; XAV939 was removed from the medium after the fifth day. Aggregates were then fed with DMEM/F12 (Gibco) supplemented with N2 (1%, R&D Systems, AR009), Glutamax (1%, Gibco), chemically-defined lipid concentrate (1%, Gibco), heparin (1 µg/mL, Sigma), and penicillin/streptomycin every other day for four days. Organoids were then embedded in Matrigel droplets and transferred to ultralow attachment six-well plates (Nunclon) in 1:1 DMEM/F12:Neurobasal medium (Gibco) with chemically-defined lipid concentrate (1%), N2 supplement (0.5%), MEM NEAA (0.5%), Glutamax (1%), beta-mercaptoethanol, N21 without vitamin A (1%, R&D Systems AR012), insulin (2 µg/mL, Gibco), and penicillin/streptomycin. Organoids received this medium every other day for seven days, transferring to an orbital shaker on the fifth day (16 days after formation). After this point, the N21 supplement was replaced with N21 with vitamin A (1%, R&D Systems AR008) and organoids were fed three times per week. After three weeks in maintenance culture, organoids were gradually transitioned to BrainPhys-based medium, in which DMEM/F12/Neurobasal base medium was replaced with BrainPhys medium (StemCell Technologies). BrainPhys-based medium was introduced in 25% increments into the DMEM/F12/Neurobasal base medium over the course of four feedings. BrainPhys medium was originally optimized for monolayer culture[94] and contains 2.5 mM glucose, a little more than 10% of the concentration in a 1:1 mixture of DMEM/F12:Neurobasal and similar to or lower than human cerebrospinal fluid. The glucose concentration in our BrainPhys-based medium was supplemented to 10 mM (ThermoFisher, A2494001), just under half the concentration of DMEM/F12/Neurobasal-based medium. Organoids were maintained for four additional weeks in BrainPhys-based medium before sample collection for a total of eight weeks of maturation.

### Low- and high-oxygen treatment

Organoids were collected in two batches. One week prior to sample collection, organoids were adapted to 10% oxygen (5% CO_2_, nitrogen balance) in a HeraCell 150i incubator (ThermoFisher). During this period, medium was equilibrated to 10% oxygen prior to feeding, and organoids were fed 24 hours before oxygen stress treatment. At the start of the experiment, plates of organoids (6-8 organoids per iPSC line per condition) were transferred to incubators maintained at 1% oxygen (5% CO_2_, nitrogen balance) or room oxygen (5% CO_2_) or left at control conditions. Oxygen concentrations were verified with a probe-based oxygen meter (Apogee Instruments, MO-200), and rapid equilibration of cell culture medium to ambient oxygen levels was confirmed in separate pilot experiments using a PreSens Fibox3 dissolved oxygen measurement device. After 24 hours, organoids were collected for single-cell dissociation.

### Single-cell RNA-sequencing sample preparation and processing

Organoids were processed using a combination of enzymatic and mechanical dissociation. Organoid medium was replaced with 1 mL papain solution (20 U/mL in EBSS, Worthington LK003150) with DNase I (100 U/mL, Worthington) supplemented with actinomycin D (5 µg/mL, Sigma A9415) and TTX (1 µM, Tocris 1069) and organoids were rapidly sheared with a pair of needles. Enzymatic digestion proceeded in the incubator, with continuous shaking, for 30 minutes. Organoids were pipetted twice with a 7-8 mm fire-polished Pasteur pipet, then returned to the incubator for an additional 10 minutes of enzymatic digestion. Samples were gently triturated four times each with fire-polished Pasteur pipets of decreasing widths (8, 6, and 3 mm) and heavy debris was allowed to settle. Samples were transferred to tubes with 2 mL inhibitor solution (3.75 mg/mL ovomucoid, 3.75 mg/mL albumin, 100 U/mL DNaseI, Worthington LK003150) and spun for 5 minutes at 200 g. Pellets were resuspended in cold Neurobasal medium with 0.5% BSA and actinomycin D and counted using a Countess II automated cell counter (Thermo) with Trypan blue. Cells from different individuals were pooled to equal concentrations, yielding three combined samples (control, low-oxygen, high-oxygen), spun down, resuspended in cold Neurobasal medium with actinomycin D, filtered with a 40 µm filter (Flowmi), and counted using a hemacytometer. Samples were loaded onto a 10x HT chip (10x Genomics) for single-cell encapsulation according to the manufacturer’s instructions, targeting approximately 3,000 cells per individual per treatment condition. Sequencing libraries were prepared using the 10x Genomics 3’ HT kit v3.1, according to the manufacturer’s instructions, in a single batch, and libraries were sequenced according to 10x Genomics specifications, targeting a minimum of 20,000 reads per cell, on an Illumina NovaSeq 6000 instrument at the University of Chicago Genomics Core Facility (RRID:SCR_019196).

### Single-cell RNA-sequencing data processing and annotation

Sequencing data were processed using the *cellranger* pipeline (v7.0.0) for read alignment (GRCh38) and cell detection. Samples were demultiplexed using *Vireo* [95] with imputed genotype information from the HapMap Project and 1000 Genomes Project, and droplets assigned to multiple individuals or with low-confidence assignments (singlet probability <0.95) were excluded. All subsequent cell filtering and annotation was performed using *Seurat* (v4.4.0) [96]. Cells were further filtered to exclude those with greater than 10% mitochondrial read content, fewer than 2,500 UMIs, or more than 20,000 UMIs, resulting in 170,841 retained cells (control = 52,671; hypoxia=57,788; hyperoxia=60,382). Data were normalized using *SCTransform* and integrated across batch and individual using *Harmony* [97] to minimize inter-individual differences in cell type annotation. All subsequent analyses made use of UMI counts, rather than transformed or fitted expression values.

Cells were annotated using a combination of reference mapping and clustering. First, cells were mapped to two published fetal single-cell datasets [52,53] using the *MapQuery* function in *Seurat*, excluding cell types absent in our organoids (microglia, endothelium, pericytes, erythrocytes), to obtain initial annotations for each cell. While most organoid cell types exhibited reasonably high concordance with fetal cell types, certain early and transitional cells could not be unambiguously annotated using the fetal reference data. To retain the information provided by these cells, which may be particularly valuable in the context of our *in vitro* experimental framework, a secondary unsupervised clustering approach was used. Dimensionality reduction and clustering were performed using *Seurat*, excluding cell cycle genes (*cc.genes* in *Seurat*) from the variable feature set to avoid spuriously co-clustering dividing radial glia and intermediate progenitor cells. A high clustering resolution was selected which produced more clusters than the fetal references and beyond which further increases did not yield interpretable changes in clusters (resolution 0.6). Clusters were then annotated based on the consensus of the fetal reference assignments of their constituent cells, or, in the case of discrepancy, additional marker gene expression, resulting in 20 cell type classes, including both “principal” cell types of the developing cerebral cortex and subtypes of neurons with regional or neuropeptide expression signatures. As fine cell type classification risks insufficient cell numbers in each group for some downstream analyses, a secondary, coarser set of annotations was created by grouping similar cell types (e.g., different inhibitory neuron subtypes) into 10 coarser classes (**Figure S1d**).

Changes in cell type abundance in response to oxygen manipulation were assessed using *propeller* [54]. Linear mixed models were estimated for both fine- and coarse-level annotations, using the treatment condition as a predictor and the parental iPSC line as a blocking variable. To further characterize sources of variation in cell type composition, additional experimental factors (collection batch, sex, iPSC passage number), were included in the model, although we note that only two iPSC lines allow direct comparisons across batches by repeated measures.

### Differential expression analysis

To identify genes differentially expressed between treatment conditions, we relied on well-established methods for analyzing bulk RNA-sequencing data. Single-cell transcriptomes were summed to pseudobulk samples, each of which corresponded to one combination of individual, treatment condition, cell type, and collection batch. Oligodendrocytes and midbrain dopaminergic neurons excluded from analysis for lack of sufficient sample sizes.

Before fitting expression models, principal component analysis was used to identify important covariates contributing to gene expression variation. Sample variation was strongly driven by the number of cells contributing to a pseudobulk sample up to 10 cells/pseudobulk sample, with a weaker effect persisting up to 20 cells/pseudobulk sample. For differential expression testing, pseudobulk samples derived from fewer than 20 cells were excluded. Pseudobulk data were TMM-normalized and genes were filtered using the *filterByExpr* function in the *edgeR* package, using treatment condition as the primary comparison group. A separate linear mixed model was estimated for each cell type using *dream* [55], with treatment condition modeled as a fixed effect and batch and parental iPSC line as random effects.

For assessing the contribution of stress-responsive cells to differential expression results, we re-ran our analysis on two datasets. In the first, we removed stress-annotated cells from each cell type before running differential expression analysis (see below). As removing cells will decrease power to detect differential expression, we generated a second dataset in which we randomly removed an equivalent number of cells of each cell type as in our stress-censored data. We compared the ratio of differentially expressed genes in the two datasets in each cell type as a measure of the transcriptional response driven by cells identified as stressed.

Accurately assessing patterns of gene regulation shared across different cell types and contexts is complicated by incomplete power. In order to characterize patterns of sharing across different cell types and treatments, we used the *mashr* package [56] to estimate posterior effect sizes and significance, with the *udr* package used in place of the default extreme deconvolution algorithm [https://stephenslab.github.io/udr/index.html]. Genes were considered significantly differentially expressed with a posterior local false sign rate (lfsr) less than 0.05, and differential expression effects were considered shared if their posterior log fold-change estimates were within a factor of 2.5 of each other.

### Enrichment analysis of DE genes

Differentially expressed genes were analyzed for enrichment of functionally defined categories using the *fgsea* package [98]. For each differential expression posterior mean and standard deviation estimate obtained from *mash*, a t-statistic was calculated and used as the ranking statistic for fgsea. The Hallmark gene sets [99] were obtained from MSigDB and used as the test set of pathways for all cell types and treatment comparisons.

### Stressed cell identification

Cells were classified as stressed by adapting the *Gruffi* framework [61], which scores local neighborhoods of cells using positive- and negative-selection gene lists. The default *Gruffi* gene lists include gene ontology terms for glycolysis and endoplasmic reticulum stress. The ER stress score did not correlate with hypoxic treatment in our data, and neither score correlated with high-oxygen exposure. As an alternative, custom gene lists were identified by *mash* as being upregulated by treatment across all fine-classified cell types (minimum two-fold change for low-oxygen treatment, minimum 1.5-fold change for high-oxygen treatment), with similar responsiveness to treatment across cell types (maximum log fold-change no more than 5 times the median log fold-change), yielding a hypoxia treatment score based on 66 genes and a hyperoxia treatment score based on 7 non-overlapping genes (**Table S3**). These genes all have known roles in stress response, redox handling, or the HIF pathway. After cell neighborhood scoring, classification was performed using *Gruffi*, using the custom gene lists as positive selection features and the default negative selection (gliogenesis-related genes) gene list. Cells were classified as hypoxia-responsive, hyperoxia-responsive, or as double-responsive, meaning they were classified by *Gruffi* as “stressed” using both gene lists. Only 223 cells were characterized as “double-responsive,” all of which were annotated as VLMC. Sensitivity to treatment was calculated as the fractional change in the proportion of any cell type classified as responsive compared to the normoxia condition.

### Immunofluorescent labeling and imaging

Organoids were washed with cold PBS and fixed with 4% paraformaldehyde (Electron Microscopy Sciences) in PBS at 4°C for 45 minutes. Organoids were then washed three times with cold PBS and cryoprotected overnight in a 30% sucrose solution before being snap frozen in OCT (Fisher). Serial cryosections (14 µm) were collected in replicate slide sets spanning the thickness of the organoid. Sections were washed with PBS and blocked with 10% NDS (Jackson) and 0.3% Triton X-100 (Sigma) in PBS for one hour at room temperature. Antibodies were diluted as follows in PBS with 2% NDS and 0.1% Triton X-100 for staining overnight at 4°C: HOPX (rabbit, 1:500, Proteintech, RRID AB_10693525), S100B (guinea pig, 1:500, Synaptic Systems, RRID AB_2620025), MKI67 (mouse, 1:500, Cell Signaling RRID AB_2797703), RELN (mouse, 3 µg/mL, DSHB RRID AB_1157892), GABA (rabbit, 1:500, GeneTex RRID AB_11173015), EOMES (rabbit, 1:500, GeneTex RRID AB_2887210), BCL11B (rat, 1:200, BioLegend RRID AB_10896795), SATB2 (mouse, 1:100, Fitzgerald AB_10809039), SOX2 (rabbit, 1:500, Synaptic Systems RRID AB_2620099), NES (mouse, 1:500, Santa Cruz RRID AB_1126569), GFAP (mouse, 3 µg/mL, DSHB N206A/8). Sections were washed four times with PBS-T (PBS with 0.05% Tween-20) and once with PBS, then incubated for two hours at room temperature with donkey secondary antibodies diluted in PBS with 2% NDS and 0.1% Triton X-100 as follows: anti-rabbit Alexa Fluor 488 (1:500, Invitrogen, RRID AB_141607), anti-guinea pig Alexa Fluor 647 (1:300, Jackson, RRID AB_2340476), anti-mouse Cy3 (1:300, Jackson, RRID AB_2340813), anti-rat Alexa Fluor 647 (1:300, Jackson, RRID AB_2340694), anti-mouse Alexa Fluor 647 (1:300, Jackson, RRID AB_2340862). Sections were washed four times in PBS-T, rinsed with water, and mounted with Fluoromount G with DAPI (Invitrogen). Slides were imaged on an Olympus VS200 Research Slide Scanner with a Hamamatsu ORca-Fusion Camera at the University of Chicago Integrated Light Microscopy Core facility, using the DAPI channel for focal mapping.

### Image analysis

For each series of organoid cryosections, the largest section was considered to be the most medial and was retained for further analysis. Image segmentation and intensity measurements were performed using the *QuPath* (v0.4.3) software package with the *Stardist* extension [100,101]. The perimeter of each section was defined using a custom pixel classifier. Cells were detected using the *Stardist* fluorescence cell detection script (dsb2018_heavy_augment.pb), with detection threshold and resolution (*pixelSize*) parameters changed (from default 0.5 to 0.3) to better suit our images. For each identified nucleus, we obtained mean fluorescence intensities and the linear distance to the nearest section edge. Cells were classified as positive for each antibody marker using a per-section threshold (1-2 standard deviations above the mean across all nuclei within the section) determined for each antibody channel. Cell types were defined conservatively from antibody markers as follows: “dividing progenitors” were defined as MKI67^+^/S100B^-^/HOPX^-^; “radial glia” were defined as HOPX^+^/S100B^-^, MKI67^+^/S100B^-^/HOPX^+^, or SOX2^+^/S100B^+^/NES^-^; “Cajal-Retzius cells” were defined as RELN^+^/GABA^-^/BCL11B^-^; “intermediate progenitors” were defined as EOMES^+^/SATB2^-^ /BCL11B^-^; “immature excitatory neurons” were defined as BCL11B^+^/SATB2^-^; “mature excitatory neurons” were defined as SATB2^+^; “inhibitory neurons” were defined as GABA^+^ (including GABA^+^/RELN^+^ and GABA^+^/BCL11B^+^); “glia” were defined as S100B^+^/HOPX^-^/MKI67^-^, S100B^+^/SOX2^+^/NES^+^, or GFAP^+^, consistent with marker combinations seen in our transcriptomic dataset. Note that these markers do not fully label all cells within a given cell type, and do not collectively cover all cell types observed by single-cell RNA-seq, but instead were chosen to localize identifiable groups of cells.

### Topic modeling

Topic modeling offers an alternative to fixed-category classification of cell states, allowing for cells to be described quantitatively by multiple gene expression programs. To alleviate the computational burden of topic model estimation and downstream analysis, single-cell data were first aggregated into 10,707 “pseudocells” by first clustering at high resolution (resolution=20) and then splitting each cluster of related cells by parental cell line and treatment condition. The *fastTopics* package [102,103] was used to fit models with a range of topics (k=10-40), and the most parsimonious model that still retained a clear hypoxia-associated topic was selected. Models were estimated by Poisson non-negative matrix factorization (*fit_poisson_nmf*) of the pseudocell count data using 400 expectation-maximization steps, followed by 200 stochastic coordinate descent steps. A multinomial topic model was obtained using the *poisson2multinom* command. Individual topics were analyzed by grade-of-membership differential expression analysis [104], comparing topic-specific DE results to cell type- or treatment-specific marker genes.

### Cis eQTL analysis

*Cis* eQTLs were identified separately for each combination of cell type and oxygen treatment condition using methods originally developed for bulk RNA-seq analysis. We obtained pseudobulk expression measurements by summing UMI counts for all protein-coding genes across cells grouped by parental iPSC line, treatment, and cell type, excluding pseudobulk samples derived from fewer than 20 cells. Cell types present in fewer than 7 individuals after pseudobulk filtering were excluded from subsequent analysis. Samples for each combination of cell type and oxygen exposure condition were TMM-normalized and expressed as log CPM values using the *edgeR* package [105]. Genes were filtered using the *filterByExpr* function in the *edgeR* package, with parameters min.count.cpm=6, min.prop.expr=0.5, and min.total.count=30, and the bottom quartile of genes ranked by standard deviation was omitted. Expression values were centered and scaled across individuals for each gene and, for each gene, rank-normalized across individuals [106]. QTL testing was performed using *MatrixEQTL* [107]. Genotype data were filtered to include variants with minor allele frequencies greater than 0.1 and Hardy-Weinberg equilibrium p-values greater than 10^-6^ using *vcftools*, and all variants within 50 kb of a gene’s transcription start site were tested for association. Gene expression principal components, obtained using the *prcomp* function in R, were used as covariates. The number of gene expression principal component covariates was chosen for each cell type and treatment so as to explain more variance in our data than in a random permutation of the data [108].

Most genetic effects on gene expression are expected to be shared across conditions. To increase our ability to detect subtle eQTL effects, *mash* was used to compare the strongest variant-gene associations across treatment conditions independently within each cell type. Note that this approach does not allow us to make rigorous statements about sharing across different cell types, but rather across treatment conditions within a single cell type. *MatrixEQTL* output was reformatted, and input data structures were created using the *fastqtl2mash* tool [56]. Because samples from different treatment conditions derive from the same parental cell lines, we the correlation structure was first estimated using *mashr*’s expectation-maximization tool. Posterior eQTL effects were considered shared across two (or more) conditions if the variant-gene pair was significant (i.e., local false sign rate < 0.05) in at least one of the conditions and the posterior effect size estimates differed by a factor of less than 2.5. Conversely, eQTL effects were considered oxygen-responsive if they were not shared in at least one oxygen condition.

### Topic-interacting cis eQTLs

Cis eQTLs that interact with cellular context were identified using *CellRegMap* [66]. The cellular environment for each pseudocell was defined by the 15-topic model estimated using *fastTopics*. Genetic similarity among pseudocells was obtained from *Plink* [109], and normalized gene expression counts for each pseudocell were used as input for *CellRegMap*. Because of the substantial computational cost of a genome-wide scan for interaction effects, tests were restricted to eGenes initially identified in the standard pseudobulk eQTL framework, testing SNPs with equal or stronger evidence of association as the *mash* lead SNP in any condition. Significant *CellRegMap* results were defined by applying a q-value threshold of 0.1 to the Bonferroni-corrected p-values.

### Comparison to disease genes

Disease-associated genes and variants were obtained from various sources (see **Tables S5 and S6**). For results obtained from the GWAS Catalog [110], intergenic variants were assigned to the gene reported in the initial study or, when no gene was reported, to the nearest gene. Traits were filtered to include at least 15 associations and no more than 500 associations, and associations were further filtered to consider only genome-wide significant results (p<5×10^-8^). Of the resulting 2989 traits, 402 were categorized as having neurological or psychiatric relevance and the remaining were considered to be “off-target” traits (**Table S6**). For gene-level analyses, we compared eGenes (genes with significant eQTL effects after *mash*) to trait-associated genes. For variant-level comparisons, we used the lead variant used by *mash*, which is the variant with the strongest association in any of the three treatment conditions used as input.

For comparisons with genes harboring rare variants, we obtained gene lists from the SCHEMA [111], SFARI (syndromic and category 1 genes) [79], Epi25 [112], BipEx [74], and Deciphering Developmental Disorders [113] projects. We filtered these gene lists using the measures of significance available for each dataset to obtain a list of 1,672 unique genes (**Table S5**).

For comparisons with GTEx eGenes, we combined the eGenes found in “Brain Cortex” and “Brain Frontal Cortex” tissue in the GTEx v8 data release. For wider comparisons to assess the novelty and tissue distribution of example eQTLs, we queried the OpenTargets Genetics [71,72] database and results from the CommonMind Consortium [73].

## Data and code availability

Sequencing data have been deposited in GEO through series accession code GSE273907. Code used to generate the results and figures in this publication are available on Github: https://github.com/bumans/organoid_oxygen_eqtl.

## Acknowledgements

We thank Natalia Gonzalez, Katherine Rhodes, and Bradley Wierbowski for manuscript comments and Josh Popp, Wenhe Lin, Peter Carbonetto, Benjamin Fair, and Yunqi Yang for technical advice. Jonathan Burnett and Olivia Allen assisted with cell culture and cryosectioning. Imaging was performed at the University of Chicago Integrated Light Microscopy Core (RRID: SCR_019197), and sequencing was performed by the University of Chicago Genomics Facility (RRID: SCR_019196). This work was supported by a grant from the National Institutes of Health (R35GM131726 to YG).

Data were generated as part of the CommonMind Consortium supported by funding from Takeda Pharmaceuticals Company Limited, F. Hoffmann-La Roche Ltd and NIH grants R01MH085542, R01MH093725, P50MH066392, P50MH080405, R01MH097276, RO1-MH-075916, P50M096891, P50MH084053S1, R37MH057881, AG02219, AG05138, MH06692, R01MH110921, R01MH109677, R01MH109897, U01MH103392, and contract HHSN271201300031C through IRP NIMH. Brain tissue for the study was obtained from the following brain bank collections: the Mount Sinai NIH Brain and Tissue Repository, the University of Pennsylvania Alzheimer’s Disease Core Center, the University of Pittsburgh NeuroBioBank and Brain and Tissue Repositories, and the NIMH Human Brain Collection Core. CMC Leadership: Panos Roussos, Joseph Buxbaum, Andrew Chess, Schahram Akbarian, Vahram Haroutunian (Icahn School of Medicine at Mount Sinai), Bernie Devlin, David Lewis (University of Pittsburgh), Raquel Gur, Chang-Gyu Hahn (University of Pennsylvania), Enrico Domenici (University of Trento), Mette A. Peters, Solveig Sieberts (Sage Bionetworks), Thomas Lehner, Stefano Marenco, Barbara K. Lipska (NIMH)

“The results published here are in part based on data obtained from the AD Knowledge Portal (https://adknowledgeportal.org). Study data were provided by the Rush Alzheimer’s Disease Center, Rush University Medical Center, Chicago, where data collection was supported through funding by NIA grants P30AG10161, R01AG15819, R01AG17917, R01AG36836, R01AG48015, U01AG46152, the Illinois Department of Public Health (ROSMAP), and the Translational Genomics Research Institute (genomic). The Mayo Clinic Alzheimers Disease Genetic Studies, led by Dr. Nilufer Taner and Dr. Steven G. Younkin, Mayo Clinic, Jacksonville, FL where data collection was supported through funding by NIA grants P50 AG016574, R01 AG032990, U01 AG046139, R01 AG018023, U01 AG006576, U01 AG006786, R01 AG025711, R01 AG017216, R01 AG003949, NINDS grant R01 NS080820, CurePSP Foundation, and support from Mayo Foundation. Study data includes samples collected through the Sun Health Research Institute Brain and Body Donation Program of Sun City, Arizona. The Brain and Body Donation Program is supported by the National Institute of Neurological Disorders and Stroke (U24 NS072026 National Brain and Tissue Resource for Parkinsons Disease and Related Disorders), the National Institute on Aging (P30 AG19610 Arizona Alzheimers Disease Core Center), the Arizona Department of Health Services (contract 211002, Arizona Alzheimers Research Center), the Arizona Biomedical Research Commission (contracts 4001, 0011, 05-901 and 1001 to the Arizona Parkinson’s Disease Consortium) and the Michael J. Fox Foundation for Parkinsons Research. And, the CommonMind Consortium supported by funding from Takeda Pharmaceuticals Company Limited, F. Hoffmann-La Roche Ltd and NIH grants R01MH085542, R01MH093725, P50MH066392, P50MH080405, R01MH097276, RO1-MH-075916, P50M096891, P50MH084053S1, R37MH057881, AG02219, AG05138, MH06692, R01MH110921, R01MH109677, R01MH109897, U01MH103392, and contract HHSN271201300031C through IRP NIMH. Brain tissue for the study was obtained from the following brain bank collections: the Mount Sinai NIH Brain and Tissue Repository, the University of Pennsylvania Alzheimer’s Disease Core Center, the University of Pittsburgh NeuroBioBank and Brain and Tissue Repositories, and the NIMH Human Brain Collection Core. CMC Leadership: Panos Roussos, Joseph Buxbaum, Andrew Chess, Schahram Akbarian, Vahram Haroutunian (Icahn School of Medicine at Mount Sinai), Bernie Devlin, David Lewis (University of Pittsburgh), Raquel Gur, Chang-Gyu Hahn (University of Pennsylvania), Enrico Domenici (University of Trento), Mette A. Peters, Solveig Sieberts (Sage Bionetworks), Thomas Lehner, Stefano Marenco, Barbara K. Lipska (NIMH)”

*The DDD study presents independent research commissioned by the Health Innovation Challenge Fund [grant number HICF-1009-003]. This study makes use of DECIPHER (*http://www.deciphergenomics.org*), which is funded by Wellcome [grant number WT223718/Z/21/Z]. See Nature PMID: 25533962 or* www.ddduk.org/access.html *for full acknowledgement*.

## Figure Legends

**Figure S1.**
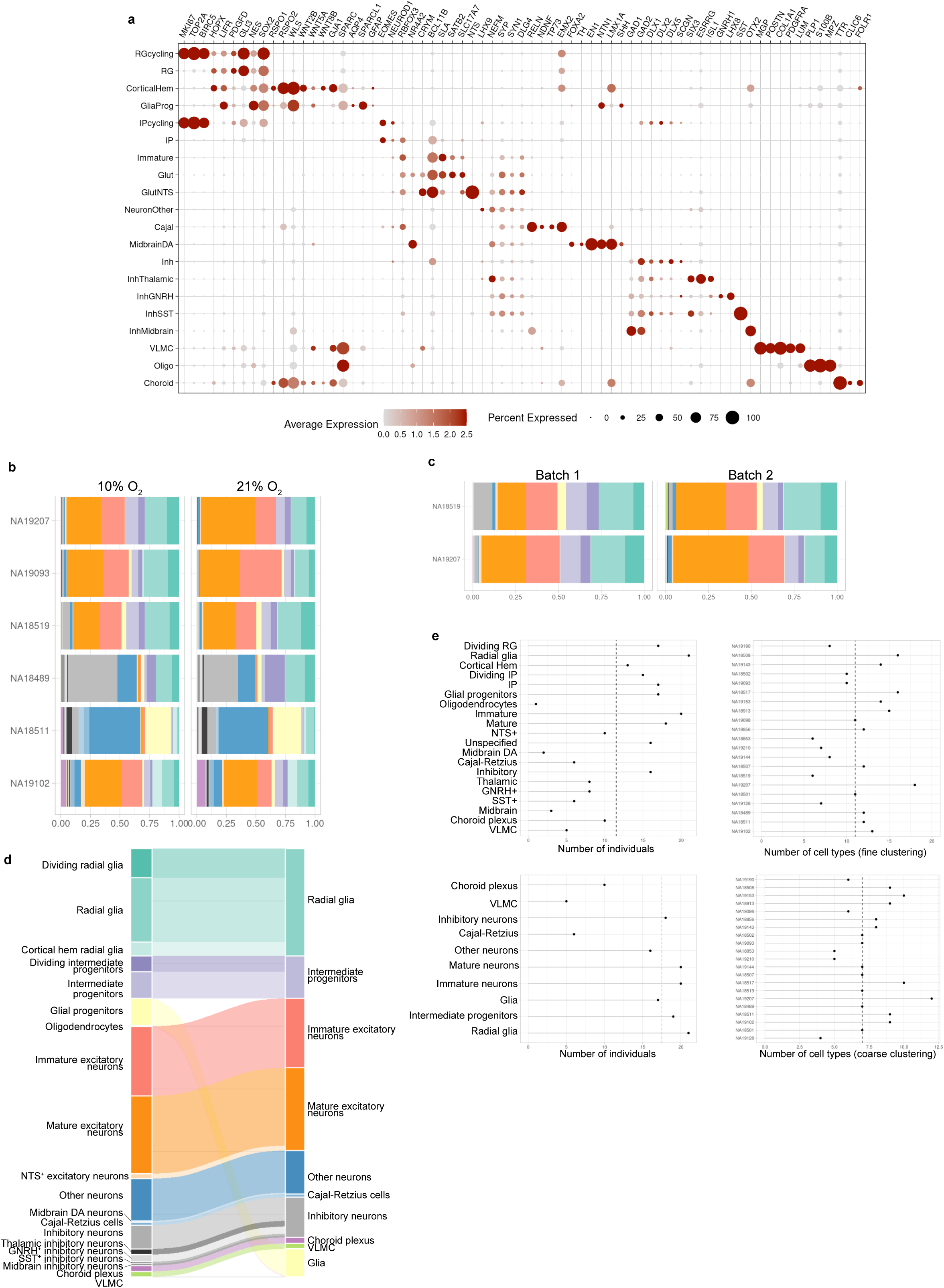
Characterization of organoid cell type composition. **(a)** Marker gene expression in each cell type annotated in Figure 1. **(b)** Organoids maintained at atmospheric (21%) or physiological (10%) oxygen for 1 week prior to collection show similar patterns of cell type composition. **(c)** Two samples were collected twice from distinct organoid formation batches. Although principal cell types are present in similar proportions, batches differ in the abundance of inhibitory neurons (gray), VLMC (lime green), mature excitatory neurons, and unclassified other neurons (blue). **(d)** Correspondence between “fine” and “coarse” classification of cell types. **(e)** Number of individuals retained per cell type after pseudobulk filtering, and number of cell types retained per individual, in the normoxia condition (representative of all sample collection conditions). Dashed lines indicate medians.

**Figure S2.**
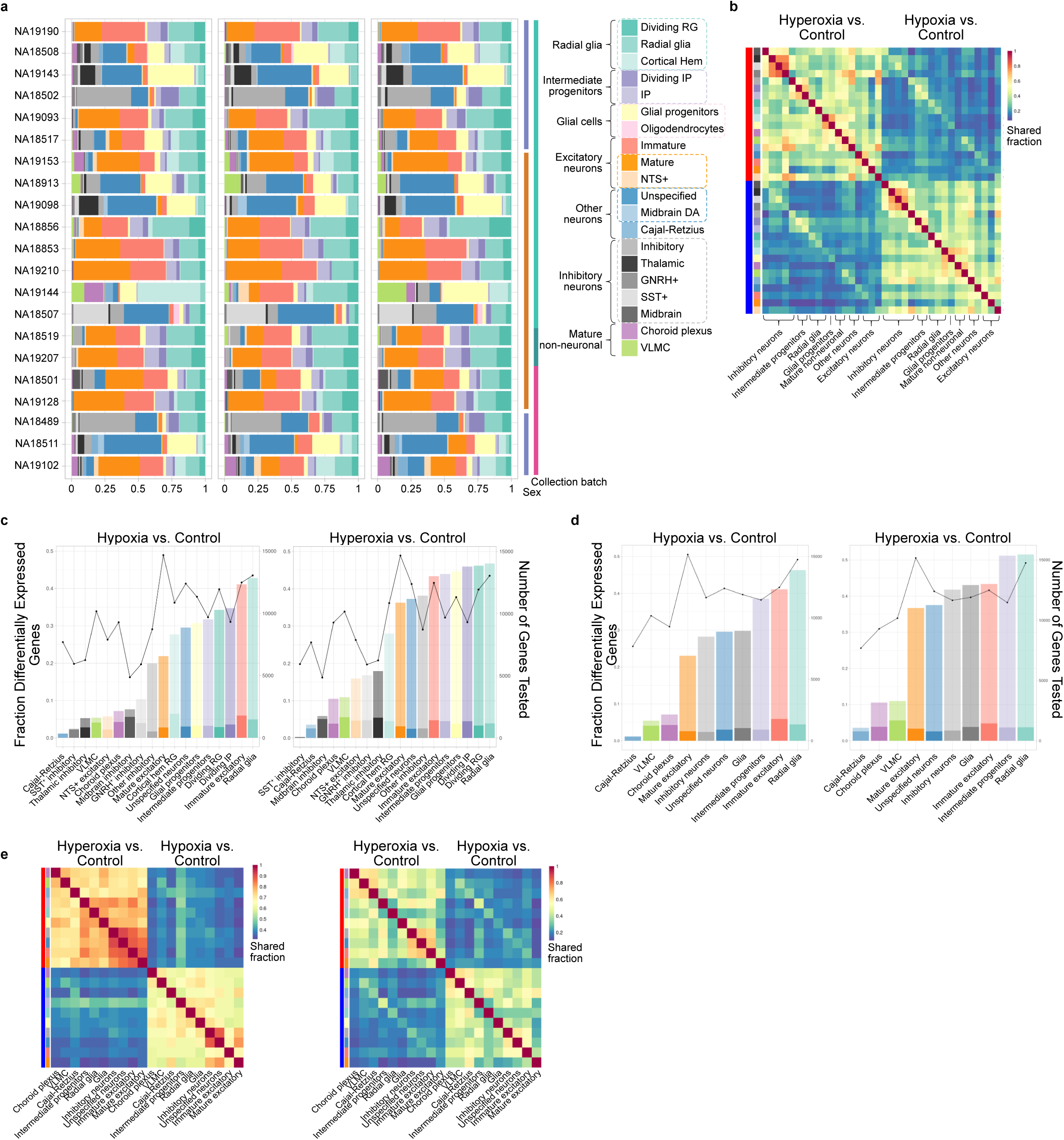
Differential expression after oxygen manipulation. **(a)** Composition of organoid samples across individuals and treatment conditions. Note that control condition data are identical to those shown in Figure 1. **(b)** Shared fraction across cell types and conditions for genes with expression changes >1.5-fold. **(c)** Proportion of tested genes in each cell type that were differentially expressed (FDR<0.05, light colors; FDR<0.05 and fold-change>1.5, dark colors). The number of genes tested in each comparison is plotted as a line (right axis). **(d)** Differential expression results for coarsely classified cells, as shown in (c). **(e)** Proportion of differential expression effects (FDR<0.05, left; FDR<0.05 and fold-change>1.5, right) shared between cell types and treatment conditions in coarsely classified cells, analogous to Figures 2a and S2b.

**Figure S3.**
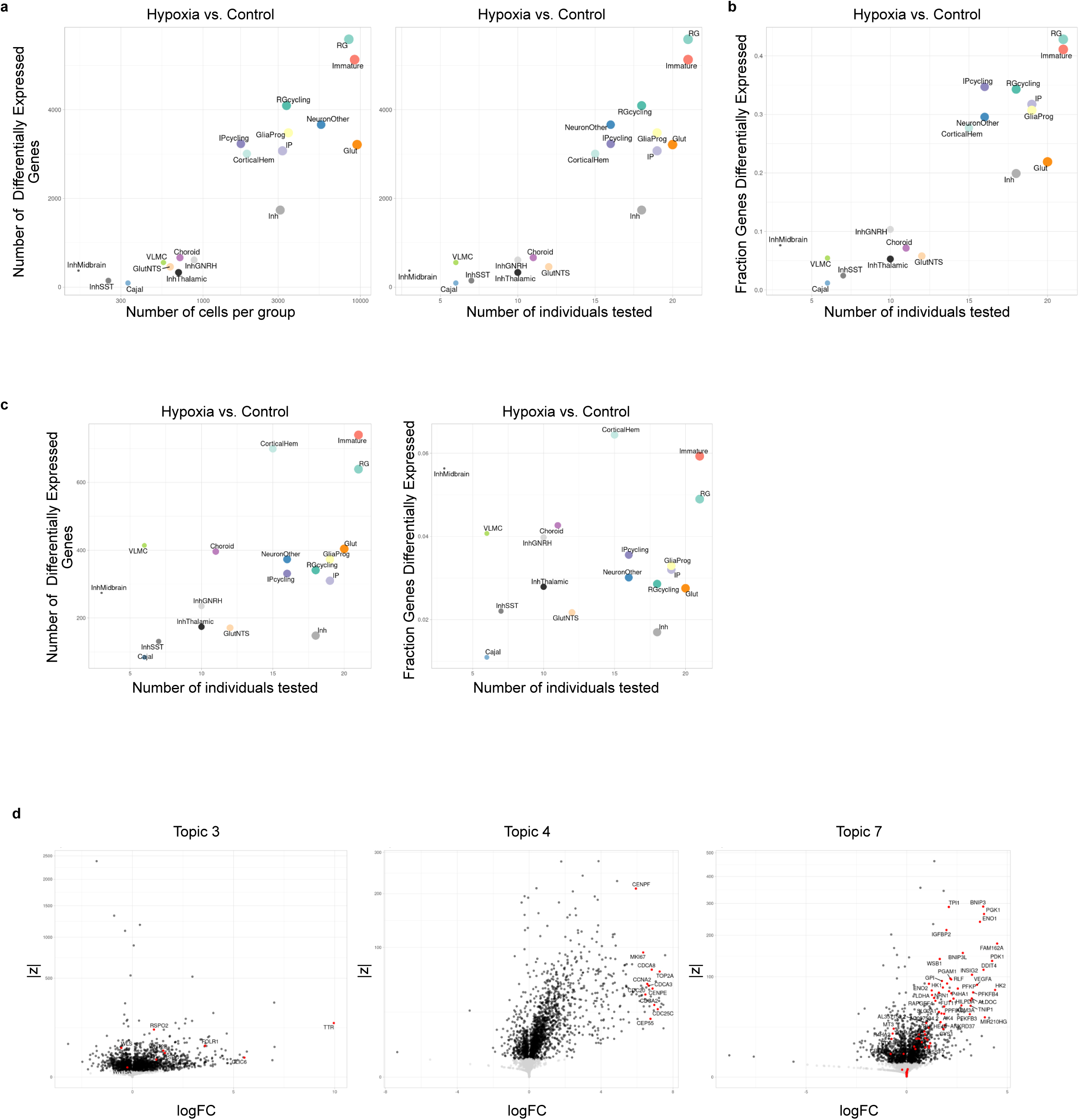
Cell abundance explains some, but not all, differences in differential expression results between cell types. **(a)** Total number of differentially expressed genes (FDR<0.05) in each cell type increases with cell type abundance, and corresponding number of individuals in differential expression comparison. **(b)** The effect shown in (a) is not driven by differences in transcriptome size or numbers of genes tested across cell types. Note that even among cell types of similar abundance, differential expression detection rate varies by almost twofold. **(c)** Results as shown in (a), excluding small differential expression effects (<1.5-fold change). Note that excess DE genes discovered in abundant cell types largely show small effect sizes. **(d)** Volcano plots of grade-of-membership differential expression testing of three topics, corresponding to Figure 2e. Choroid plexus markers are highlighted for topic 3, dividing cell markers are highlighted in topic 4, and cell type-shared hypoxia-response genes are highlighted in topic 7.

**Figure S4.**
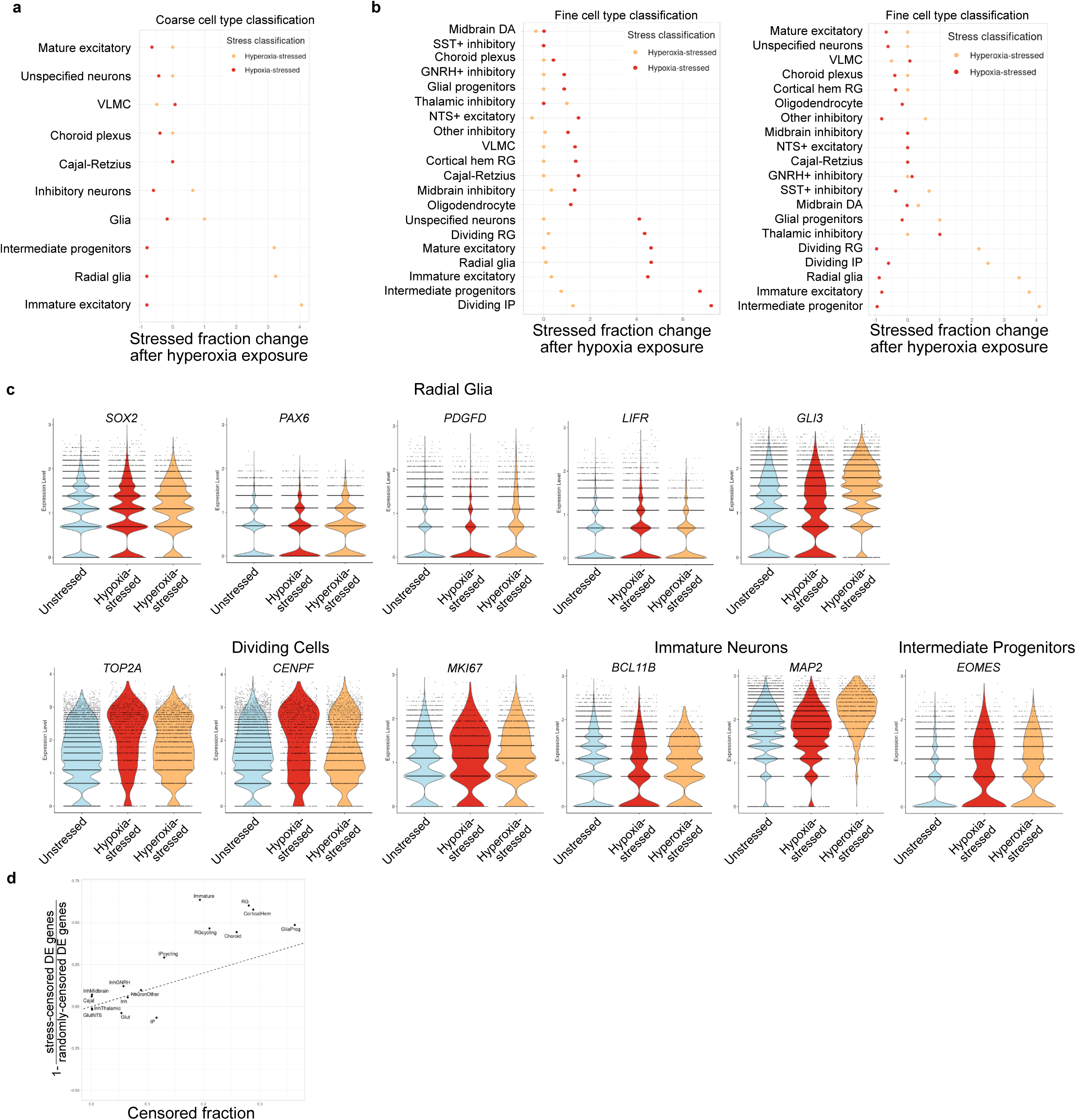
Stressed cell identification and responses to treatment. **(a)** Fractional change in cell proportions classified as hypoxia-stressed and hyperoxia-stressed after high-oxygen treatment in coarsely classified cells. Note that elevated environmental oxygen reduces the fraction of cells classified as hypoxia-stressed. **(b)** Fractional change in cell proportions classified as hypoxia-stressed and hyperoxia-stressed after low- or high-oxygen treatment using fine-grained cell type classifications. **(c)** Canonical cell type markers are maintained in stressed cells. Cells classified as “stressed” retain key markers of their identities. **(d)** Stress-responsive cells account for many, but not all, of the DE genes (FDR<0.05 and fold-change>1.5) induced by hypoxia in the most responsive cell types. Censoring stressed cells reduces DE genes more than randomly censoring matched proportions of cells for indicated cell types.

**Figure S5.**
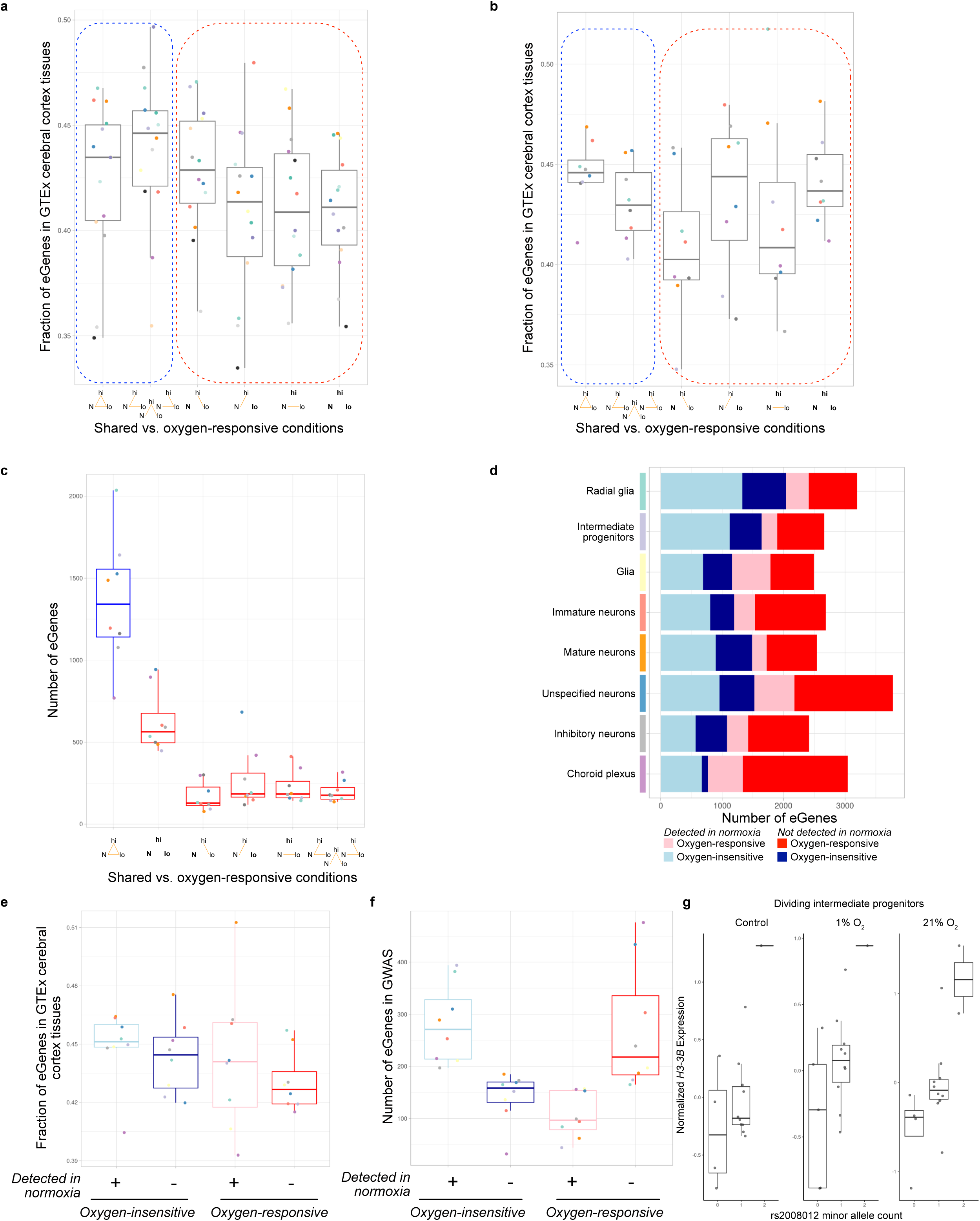
Oxygen-responsive eGenes are less abundant in GTEx and include large numbers of GWAS genes. **(a)** Fractions of eGenes in each category classified as eGenes in GTEx cerebral cortex tissues. Oxygen-responsive eGenes boxed in red are less likely to be present in GTEx compared to eGenes boxed in blue (p=0.0083, one-sided paired Wilcoxon test). **(b)** Results as shown in (a) for eGenes identified from coarsely classified cell types (p=0.0039, one-sided paired Wilcoxon test). **(c)** Number of eGenes identified in each coarsely classified cell type, analogous to Figure 3a. **(d)** Number of eGenes identified in each coarsely classified cell type, analogous to Figure 3b. **(e)** Fractions of eGenes, identified in coarsely classified cell types, in each category classified as eGenes in GTEx cerebral cortex tissues, analogous to Figure 3c (p=0.055, one-sided paired Wilcoxon test of oxygen-insensitive category [blue] against oxygen-responsive/undetected in normoxia category [dark red]). **(f)** Number of eGenes in each coarsely classified cell type which are the nearest gene to a genome-wide significant GWAS finding among 402 brain-related traits, analogous to Figure 4b. **(g)** Non-dynamic (“standard”) eQTL effect of rs2008012 in dividing intermediate progenitor cells. This eQTL has an oxygen-responsive effect in immature neurons (Figure 4c).

**Table S1.** Results from *propeller* testing of oxygen treatment, sex, collection batch, and iPSC passage number on cell type proportions.

**Table S2.** Results from differential expression testing using *dream*.

**Table S3.** Gene lists used for identifying oxygen-responsive cells.

**Table S4.** Results of eQTL mapping and *mash* analysis.

**Table S5.** Rare variant genes used for analysis of eQTL results and rare variant gene-eQTL-GWAS phenotype triads.

**Table S6.** GWAS study accession identifiers (GWAS Catalog) used for analysis of eQTL results.

